# Compartmentalized oxygen generation from hydrogen peroxide enables anaerobic oxidative phosphorylation and confers a fitness advantage

**DOI:** 10.64898/2026.08.01.742219

**Authors:** Stefan Urs Moning, Yannick Bärtschi, Lucia Heger, Jonas Hentschel, Jan Kägi, Thorsten Friedrich, Christoph von Ballmoos

## Abstract

Hydrogen peroxide is a host-derived reactive oxygen species that invading bacteria must detoxify to overcome the host defence. It was previously proposed that cytoplasmic catalase converts this peroxide into molecular oxygen, allowing facultative anaerobes such as *Escherichia coli* and *Salmonella* to respire in oxygen-limited niches. Here, we experimentally test this mechanism by reconstituting a minimal system comprising bovine catalase and purified *E. coli* quinol oxidases in liposomes. We find that catalase-generated oxygen sustains quinol oxidation under anaerobic conditions, and the rapid consumption keeps the environment effectively oxygen-free. When F_1_F_O_ ATP synthase is included into the proteoliposomes, the system generates a protonmotive force that drives ATP synthesis, with activity scaling with hydrogen peroxide concentration. Competition experiments at low peroxide concentrations demonstrate that compartmentalized oxygen production provides a decisive advantage, as liposomes lacking catalase are strongly outcompeted. All three terminal quinol oxidases of *E. coli* were tested and shown to support ATP synthesis *in vitro*, suggesting that *bd*-II involvement *in vivo* is likely due to regulation of gene expression rather than catalytic constraints. Together, our data show that intracellular oxygen generation from host-derived peroxide enables oxidative phosphorylation in globally anaerobic, electron-acceptor-limited environments such as the gut, thereby providing a mechanistic explanation for the observed fitness advantage.

## Introduction

In most organisms, the majority of the universal energy currency ATP is produced by the rotary F_1_F_O_ ATP synthase driven by a transmembrane electrochemical proton gradient generated by primary proton pumps. Depending on environment and organism, various types of protonmotive force (*pmf*)-generating systems are found, i.e. light-driven, decarboxylation-driven, and redox-driven systems [1]. The efficiency of this process determines the rate of bacterial growth and is thus elementary for competitive growth environments of high microbial density such as the human digestive tract [2,3].

Within the redox-driven systems, a large variety of enzyme combinations leading to *pmf* generation by primary proton pumps exist, where aerobic respiratory chains as found in many bacteria and mitochondria are best known and represent the most efficient mode of energy conservation [4,5]. Here, reduction equivalents such as NADH and succinate are oxidized to reduce molecular oxygen to water, and the free energy of this exergonic reaction enables redox-driven proton transport across biomembranes [6]. Oxygen is the preferred electron acceptor; however, under anoxic conditions, it is replaced by nitrate, fumarate, or sulfate [7], resulting in lower ΔG yields. The competition for the best possible acceptors can be illustrated by the example of the human gut, where anaerobic bacteria utilize exotic acceptors such as thiosulfate, trimethylamine N-oxide or DMSO to define their nutrient niche. Per definition, facultative anaerobic bacteria can thrive in both aerobic and anaerobic environments. This adaptability gives pathogens a selective advantage that enables them to invade tissues, survive host defences, and persist under varied and changing oxygen conditions often found during infection. Most life-threatening bacterial pathogens, including many on the WHO’s list of priority antibiotic-resistant threats like *Escherichia coli* and *Salmonella*, are facultative anaerobes [8–12].

At the core of aerobic *pmf* generation in *E. coli* are the three terminal quinol dependent oxidase *bo*_3_, *bd*-I and *bd*-II [4,13,14]. The genes of *bo*_3_ oxidase are expressed at high oxygen concentrations. The *bo*_3_ oxidase belongs to the family of *heme*-copper oxidases and acts as a true proton pump (2H^+^/e^-^). In contrast, the genes of the two *bd-*type oxidases are expressed at low oxygen conditions [15,16] and create a *pmf* by charge separation (1H^+^/e^-^) (i.e. uptake of cytoplasmic protons to the catalytic site close to the periplasmic side and release of protons from quinol to the periplasm) [17–19].

After infection by bacterial pathogens, host epithelial and immune cells produce reactive oxygen species such as superoxide and hydrogen peroxide (H_2_O_2_) *via* membrane bound NADPH oxidases to ward off invading bacteria as first line of defence. Crucially, it has been recognized that pathogenic *Citrobacter* and *Escherichia coli* strains have evolved strategies to exploit host derived H_2_O_2_, providing them a fitness advantage upon infection [20,21]. It was shown that the *app*BCX gene cluster encoding the *bd*-II oxidase in *E. coli* strain Nissle 1917, provides a growth advantage during colitis induced by dextran sulfate sodium in mice. This advantage was lost in mice lacking NOX1, an epithelial enzyme that produces reactive oxygen species (ROS), including H₂O₂ [20]. The authors proposed that H_2_O_2_ produced by NOX1 diffuses into bacteria where it is broken down by cytoplasmic catalases into water and oxygen. They hypothesize that the produced oxygen serves as terminal electron acceptor for aerobic respiration by *bd*-II oxidase, but not *bd*-I oxidase, thereby increasing their bioenergetic efficiency leading to the observed growth advantage [22]. In this study, we address the hypothesis of localized oxygen production by catalase in a bottom-up system, using purified ATP synthase and quinol oxidases from *E. coli* reconstituted into liposomes. Competition experiments between liposomes containing or lacking catalase are used to quantify the importance of compartmentalized oxygen generation from hydrogen peroxide.

## Results and Discussion

### Coupling of hydrogen peroxide breakdown to quinol regeneration

The majority of our experiments were performed using *E. coli bd*-II oxidase that was purified by metal affinity chromatography and gel filtration following described protocols [18]. To mimic the activity of cytoplasmic catalase, we used commercially available catalase from bovine liver [23]. First we aimed to find out, whether this catalase stoichiometrically forms oxygen from hydrogen peroxide, and whether this oxygen is a substrate for purified *bd*-II oxidase supplied with pre-reduced quinol (Figure 1A).

**Figure 1:**
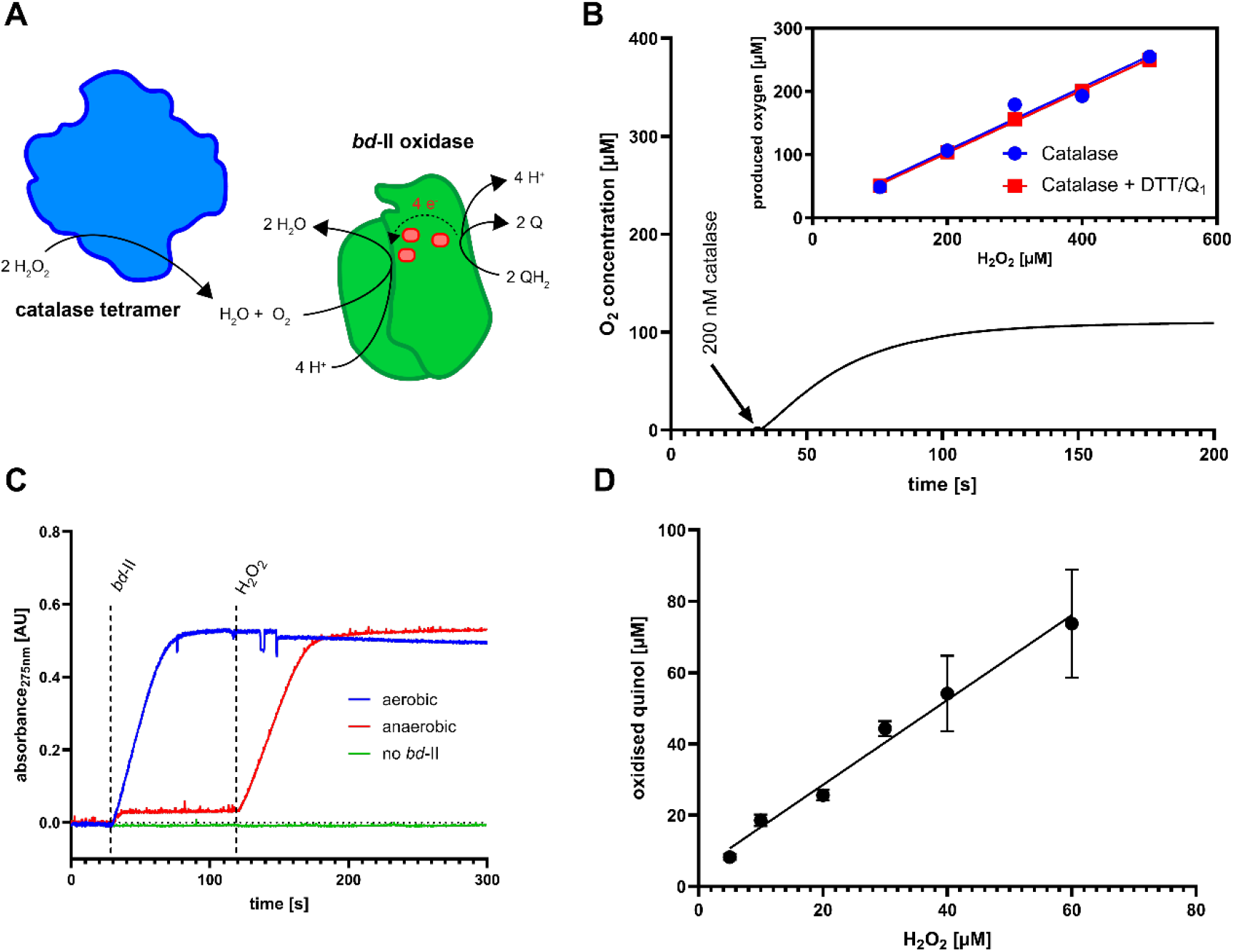
Oxidation of quinol initiated by hydrogen peroxide by bovine liver catalase and. E¡.coli.bd**-II oxidase.** A) Schematic representation of all reactions involved in hydrogen peroxide driven quinol oxidation. Bovine liver catalase (blue) dissociates hydrogen peroxide into water and oxygen. *bd*-II oxidase from *E. coli* (green oxidizes quinol (QH_2_) to quinone (Q) and transports the electrons over three heme groups to molecular oxygen, which is reduced to water. B) Production of oxygen by bovine liver catalase measured using a Clark-type electrode. A baseline corrected raw measurement is shown where 200 µM of hydrogen peroxide is processed by bovine liver catalase to molecular oxygen. Insert: Titration of hydrogen peroxide to catalase and influence Q_1_/DTT on oxygen production. C) Quinol oxidation by *bd*-II oxidase measured at 275 nm. Reactions were initiated by 25 nM *bd*-II and 200 µM H_2_O_2_ with 25 nM catalase present. Anaerobic buffer was obtained by bubbling with nitrogen in an air-tight cuvette. D) Titration of the H_2_O_2_ / catalase / oxygen / *bd*-II reaction cascade at anaerobic conditions with 25 nM of *bd*-II and catalase.

The catalase activity under anaerobic conditions was assessed using an oxygraph, in which an anaerobic buffer was supplied with different amounts of hydrogen peroxide and equilibrated, before a fixed amount of purified catalase was added. As depicted in Figure 1B, generation of oxygen occurred immediately and continuously until all peroxide was used up. Analysis of oxygen generation at different peroxide concentration showed a strong linear dependency with a O_2_/H_2_O_2_ stoichiometry close to expected value of 0.5 (Figure 1B, inset). The reaction was not disturbed by the short chain ubiquinone Q_1_ and the reducing agent DTT, which was used in later experiments. Next, we verified that the catalase-derived oxygen is consumed by purified *bd*-II oxidase in the presence of pre-reduced quinol serving as electron donor. The activity was measured by following the absorbance at 275 nm, where oxidation of quinol to quinone is observed. As depicted in Figure 1C, addition of purified *bd*-II oxidase to anaerobic buffer containing quinol yielded only a minor absorption increase, most likely due to the small oxygen amount in the enzyme sample. Upon addition of peroxide, quinol oxidation is readily observed at similar rates as under aerobic conditions, indicating that not the catalase but the oxidase reaction is rate limiting. No spontaneous oxidation of quinol was observed in the absence of *bd*- II oxidase but presence of hydrogen peroxide, confirming that the reaction was exclusively enzyme-mediated. Titration of this reaction (Figure 1D) shows that ∼1.2 quinol is oxidized per peroxide molecule present, slightly more than the expected ratio of 1.

### Assembly of a compartmentalized system

Next, we reconstituted *bd*-II oxidase into catalase containing liposomes, mimicking the compartmentalisation of the bacterial cytoplasm (Figure 2A). Orientation of the quinol oxidase is not relevant at this step, as short-chain quinones are membrane permeable and can reach either population. As in our previous experiments with *bd*-I oxidase [24], pre-formed liposomes were treated with non-solubilizing concentrations of sodium cholate, incubated with purified oxidase, and the excess sodium cholate was removed using gel filtration [24]. To ensure efficient encapsulation of the large catalase, the enzyme was added in the lipid rehydration buffer used to resuspend the dried lipids, followed by liposome formation *via* freeze-thawing and size homogenization using extrusion. These liposomes were used for detergent-mediated reconstitution of *bd*-II oxidase, and the non-encapsulated catalase was removed by ultracentrifugation. As depicted in Figure 2B and S1, catalase was successfully encapsulated and retained during reconstitution, and catalase activity was detected after ultracentrifugation. We titrated the activity against known amounts of soluble catalase and calculated an average number of 4 catalases per liposome. Next, we tested reconstituted *bd*-II activity in liposomes by following oxidation of pre-reduced ubiquinol Q_1_ that readily partitions into the liposomal membrane. Similar to our observation with solubilized *bd*-II oxidase under aerobic conditions (Figure 1C), quinol oxidation started immediately upon proteoliposome addition. No quinol oxidation is observed under anaerobic conditions, until hydrogen peroxide is added, which is converted to oxygen by catalase (Figure 2C). As the catalase-derived oxygen is produced within the liposomes, it is either directly used by the oxidases presenting the oxygen channel to the liposome’s interior or it has to penetrate the liposomal membrane, before it reacts with the oxidases reconstituted in the opposite orientation. The hydrophobic character of oxygen supports its accumulation in the membrane. We therefore asked, if the reconstituted *bd*-II oxidase is capable of consuming all catalase produced oxygen before it reaches the extra- liposomal phase from where it can be detected by the electrode of an oxygraph instrument. To this end, we placed an anaerobic solution containing quinol and catalase-liposomes with or without reconstituted *bd*-II oxidase into an oxygraph chamber and followed oxygen production upon hydrogen peroxide addition (Figure 2D). With this experimental setup, only extraliposomal oxygen reaches the sensor of the device. To ensure a continuous supply of electrons, we added 4 mM DTT to the solution, ensuring constant reduction of the ubiquinone pool. As depicted in Figure 2D, with liposomes containing only catalase, a linear dependence of added hydrogen peroxide and observed oxygen at the electrode was obtained. In contrast, with liposomes containing both catalase and *bd*-II oxidase, no oxygen was detected up to a concentration of 150 μM hydrogen peroxide, demonstrating that aerobic respiration takes place under otherwise anaerobic conditions. At higher concentrations, the number of reconstituted *bd*-II oxidases is not sufficient to completely block oxygen loss, although the measured concentrations remain low, supporting the idea that oxygen preferably “waits” in the membrane.

**Figure 2:**
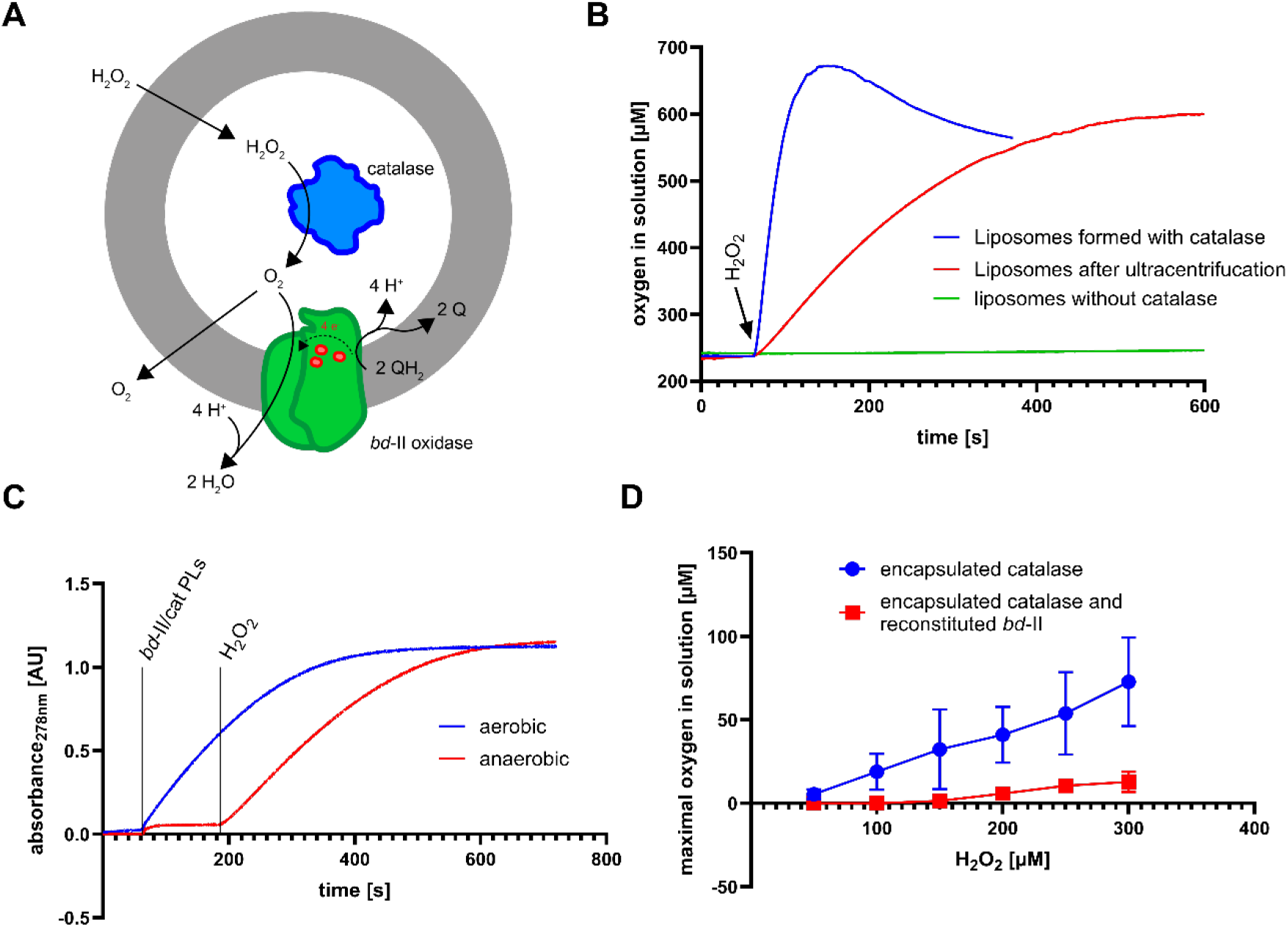
Integration of bovine liver catalase and E¡.coli bd-II oxidase into liposomes. A) Schematic representation of liposomes with *bd*-II oxidase reconstituted and bovine liver catalase encapsulated. Only one orientation is shown for simplicity, but a non-unifrom orientation is expected. Hydrogen peroxide diffuses through the lipid membrane and is processed by catalase, providing oxygen for *bd*-II oxidase. B) Hydrogen peroxide induced oxygen production of liposomes formed in presence of catalase before and after removal of exterior catalase and of liposomes formed without catalase. C) Quinol reduction by proteoliposomes containing *bd*-II oxidase and bovine liver catalase measured at 275 nm. Reactions were initiated by 0.1 mg/ml *bd*-II/catalase proteoliposomes and 200 µM H_2_O_2_. Anaerobic buffer and proteoliposomes were obtained by stirring in an anaerobic glovebox. D) Oxygen concentration in solution upon addition of hydrogen peroxide to proteoliposomes with catalase encapsulated with and without *bd*-II reconstituted. Maximal oxygen concentration in solution is plotted against the concentration of hydrogen peroxide.

### Hydrogen peroxide dependent ATP synthesis

In contrast to terminal oxidases of the heme copper superfamily like *bo*₃ oxidase and *aa*₃ oxidase, *bd* oxidases have a different arrangement of redox groups, which is not compatible with physical proton pumping across the membrane. Based on single turnover measurements and corroborated by the recent molecular structures, however, *bd* oxidases are believed to generate to a *pmf* by electrogenic uptake of protons from the cytoplasm to the membrane located catalytic site required for water formation [18,19]. Simultaneously, protons released during oxidation are proposed to leave specifically to the periplasmic side. Together, these two processes result in a net proton transfer across the membrane from the cytoplasm to the periplasm. The uptake of protons to the catalytic site is electrogenic, generating a membrane potential Δψ. A major consumer of the *pmf* and of periplasmic protons is the F_1_F_O_ ATP synthase that produces cytoplasmic ATP from ADP and phosphate by its rotatory mechanism [25–27].

Here we mimic this situation by co-reconstitution of *bd*-II oxidase and ATP synthase into liposomes containing catalase (Figure 3A), making *bd*-II the only possible source for the generation of *pmf*. All enzymes were mixed during the reconstitution process and proteoliposome formation containing bd-oxidase and and ATP synthase was verified using cryoEM (Figure S2). In the micrographs, the ATP synthase preferably orients with its F_1_ head towards the outside of the liposomes similar to what has been reported using this reconstitution method [28], which is inverted from the cellular situation. Accordingly, orientation of the *bd* oxidase must also be predominantly inverted to produce a *pmf* that is positive inside, driving protons out through the ATP synthase and producing ATP. Here, we do not control *bd* oxidase orientation deliberately, and oppositely oriented enzymes might cancel each other out.

**Figure 3:**
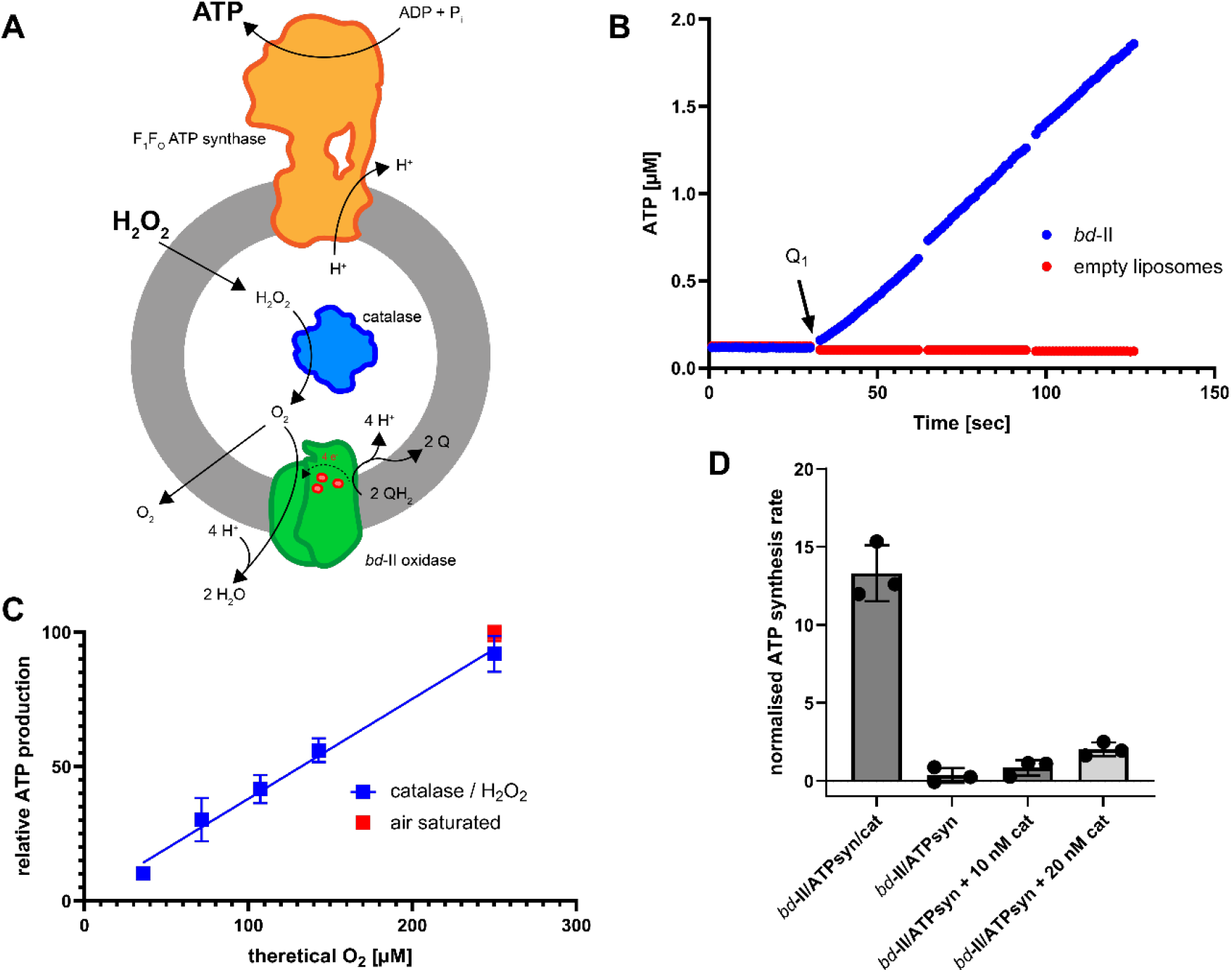
Hydrogen peroxide dependent production of ATP by F_1_F_O_ ATP synthesis. A) Schematic representation of a proteoliposome capable of ATP synthesis upon addition of hydrogen peroxide (simulation the *in vivo* process postulated by Chanin et al [20]). Hydrogen peroxide diffuses into the proteoliposome, enabling generation of oxygen by bovine liver catalase, *pmf* formation by the terminal oxidase *bd*-II from *E. coli* and finally *pmf* dependent ATP production by the *E. coli* F_1_F_O_ ATP synthase. B) ATP synthesis driven by *E. coli bd*-II oxidase by *bd*-II/ATP synthase soybean lecithin proteoliposomes at aerobic conditions with electrons delivered by 4 mM DTT and started by addition of 15 µM Q_1_. The empty liposomes control did not contain any *bd*-II oxidase. C) Titration of hydrogen peroxide to soybean lecithin proteoliposomes with *bd*-II and ATP synthase co-reconstituted and catalase encapsulated at anaerobic conditions. The reactions were run in presence of Q_1_/DTT in an anaerobic glovebox and started by addition of hydrogen peroxide. All measurements were normalised to the ATP synthesis rate under aerobic conditions (100%). D) Anaerobic ATP synthesis of soybean lecithin proteoliposomes with *bd*-II and ATP synthase co-reconstituted with encapsulated catalase, without catalase and with equal amounts of catalase in the outside buffer energised by Q_1_/DTT. 10 nM catalase has been found to be equally active as 0.1 mg/ml liposomes with catalase encapsulated. Synthesis rates were normalised to the aerobic ATP production (100%) of the corresponding proteoliposomes.

However, using a luciferin/luciferase system, we show that the system described here is indeed capable of producing ATP continuously (Figure 3B), promoting the following two conclusions.

Firstly, *E. coli bd*-II oxidase is indeed a *pmf* generating quinol:oxygen oxidoreductase and that passively delivered protons from quinol oxidation to the inside of the liposomes are sufficient to allow continuous ATP synthesis (Figure 3B). ATP synthesis is immediate and continuous, excluding rapid exhaustion of protons in the small intraliposomal volume that would lead to a rapid pH increase and prohibit ATP synthesis. Secondly, like for *bd*-I, *bo*_3_ and *aa*_3_ oxidases, this reconstitution methods produces liposomes with *bd*-II oxidases preferably in the inside-out orientation [24,29,30]. To investigate, whether ATP synthesis is also possible by catalase- derived oxygen under otherwise anaerobic conditions, we adjusted our measurement technique. To ensure a fully anoxic environment, the experiment was performed in an anaerobic glove box, and samples were taken at the indicated time points and quenched in tributyltin chloride containing buffer, a strong ATP synthase inhibitor [31]. After the experiments, samples were exported from the glovebox and mixed with luminescence assay, and the ATP was determined using standardized ATP amounts (Figure S3AB). As depicted in Figure 3C, ATP synthesis in the presence of H_2_O_2_ is indeed observed in *bd*-II/ATP synthase liposomes containing 4 molecules of catalase (∼15 μM in a 100 nm liposome), while no notable ATP production is observed in proteoliposomes lacking catalase (Figure 3D and Figure S3C). If run to completion, the total amount of ATP synthesis was directly proportional to the concentration of peroxide present, and 500 μM peroxide lead to a similar amount of ATP to that obtained when the reaction was carried out under aerobic conditions (∼250 μM O_2_) using the same analysis technique (Figure 3C). In contrast, if the same number of catalase molecules (Figure S3D) is added to the outside of proteoliposomes containing no entrapped catalase, 12-times less ATP is produced (Figure 3D). In other words, while the same amount of oxygen was produced by catalases, the oxygen produced within liposomes was ∼12-times more efficient in producing ATP. Even if the external catalase amount was doubled, the difference was more than 5-fold.

To simulate competition from other oxygen-consumers in the gut, we introduced proteoliposomes containing solely reconstituted quinol oxidase, competing for oxygen without contributing to ATP production (Figure 4A). Titration of these competing proteoliposomes against *bd*-I/ATP synthase/catalase proteoliposomes revealed no reduction in ATP synthesis following the addition of 50 µM hydrogen peroxide, even under conditions of excess competition (Figure 4B). This observation further supports that the oxygen generated inside the liposomes is utilized predominantly by the reconstituted oxidase within the catalase containing vesicles and does not diffuse out of the vesicles. In contrast, when catalase was not encapsulated but instead added to the bulk solution at equivalent amounts, the oxygen produced became accessible to both proteoliposome populations, resulting in substantial competition for the available oxygen (Figure 4C). Under these conditions, ATP production decreased to approximately 30% at a 1:1 ratio of ATP producing and competing proteoliposomes. A potential explanation for this proportionally higher reduction in ATP synthase rate might be a slightly higher reconstitution efficiency of oxidase in liposomes lacking ATP synthase. Taken together, these results demonstrate that compartmentalized oxygen production from hydrogen peroxide *via* catalase supports robust ATP synthesis, and that only internally generated oxygen can efficiently serve as an electron acceptor.

**Figure 4:**
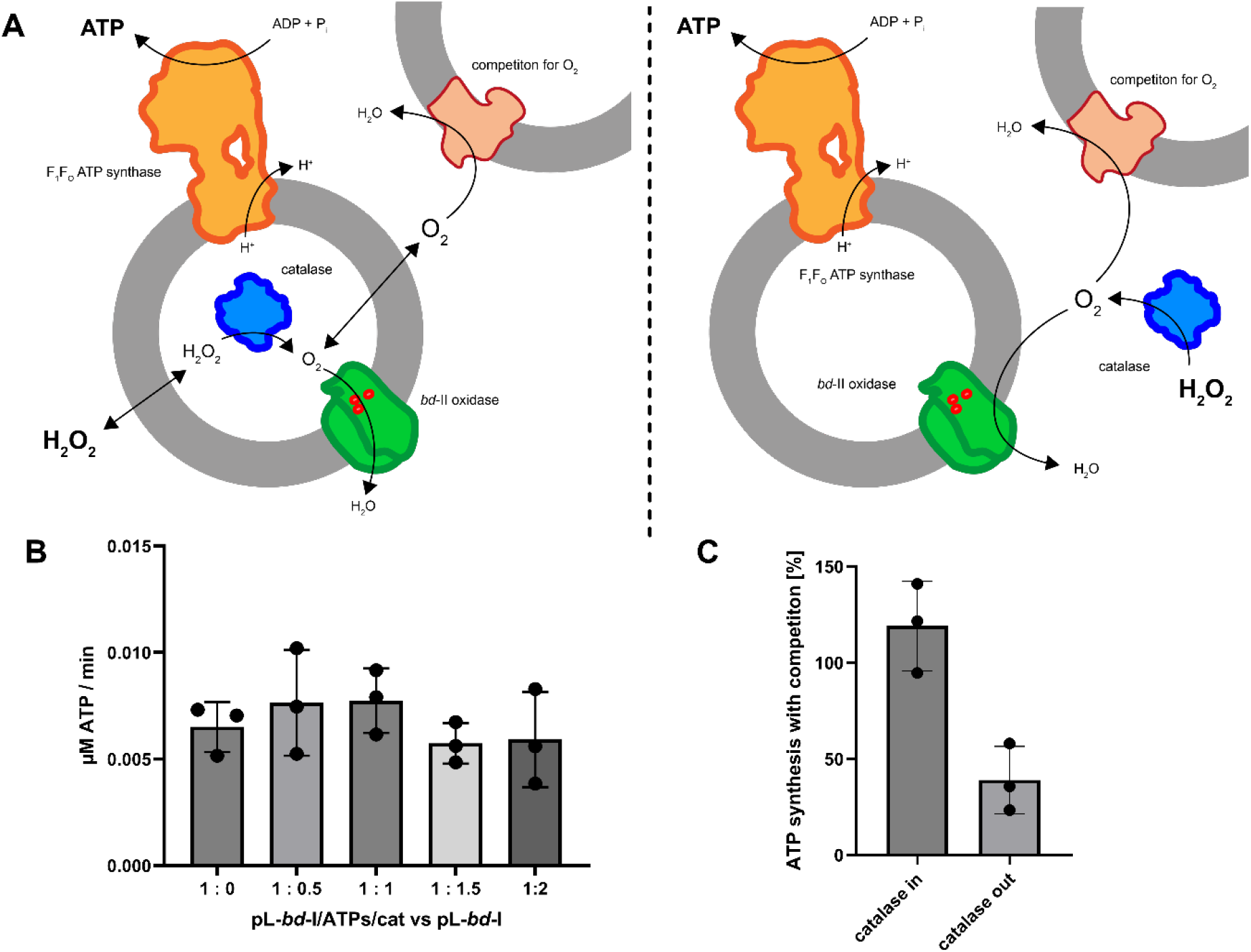
Hydrogen peroxide driven ATP synthesis with competing oxygen consumers. A) Schematic representation of minimal respiratory system (*bd*-II, F_1_F_O_ ATP synthase, catalase) with external competition (*bo*_3_ oxidase). Oxygen is produced either in the proteoliposomes by encapsulated catalase and hydrogen peroxide diffusing through the membrane (right). The produced oxygen is either used by the *bd*-II oxidase reconstituted into the same proteoliposome for pmf generation and ATP synthesis or diffuses into the bulk solution, where it can also be used by *bo*_3_ oxidase, reconstituted without ATP synthase. If oxygen is produced outside of the proteoliposomes (left) it can freely diffuse to both *bd*-II/ATP synthase and *bo*_3_ proteoliposomes, but only proteoliposomes with ATP synthase are able to produce ATP. B) Competition for oxygen produced inside the proteoliposomes. Soybean lecithin proteoliposomes with *bd*-I and ATP synthase co-reconstituted and catalase encapsulated were mixed in different ratios with proteoliposomes only containing *bd*-I oxidase, unable to produce ATP. Liposome concentration was kept constant supplementation with empty liposomes. ATP synthesis at anaerobic conditions was driven by Q_1_/DTT and started by addition of 50 µM of hydrogen peroxide. C) Competition for oxygen produced by catalase either encapsulated into the proteoliposome or added to the bulk buffer. Soybean lecithin proteoliposomes with *bd*-I and ATP synthase co-reconstituted with and without catalase encapsulated were mixed in a 1:1 ratio with proteoliposomes only containing *bd*-I oxidase, unable to produce ATP. Rates were normalised to ATP synthesis rates of liposomes without competition.

### Role of E¡.coli terminal oxidases

At the core of the findings by Chanin et al. is the observation that together with catalase, only cytochrome *bd*-II (AppBCX), but not *bd*-I (CydABX) provides a fitness advantage to *E. coli* in the presence of host-derived hydrogen peroxide during intestinal inflammation. Current models suggest this distinction is largely rooted in transcriptional regulation, as the expression of *bd*-II increases upon exposure to stressors like nitrate and H_2_O_2_ under anaerobic conditions, whereas the expression of *bd*-I does not [13,14]. Moreover, the proton-pumping *bo*_3_ oxidase is not expected to be expressed at all under anaerobic conditions. It thus remains a critical question whether intrinsic biochemical and catalytic activities beyond transcriptional differentiation contribute to the observed phenotype. In the past, in addition to oxidase activity, also non-canonical peroxidase and catalase activities were described for both *bd* type oxidases, which would allow ATP synthesis in the absence of catalase. However, reports to these activities are conflicting [32–34].

To help resolve these contradictions, we compared whether oxygen produced by catalase also serves as a substrate for the other *E. coli* quinol oxidases, *bd*-I and *bo*_3_, enabling them to create a *pmf* that drives ATP synthesis. We purified *bd*-I, *bd*-II and *bo*_3_ oxidase and measured oxygen production following the addition of 200 μM hydrogen peroxide to follow catalase-like activity (Figure S4). None of the oxidases showed significant catalase-like activity. Next, we co-reconstituted each of the three oxidases together with ATP synthase into liposomes containing catalase. As depicted in Figure 5A (red bars), the amount of ATP was in the same order of magnitude. While we used the same amount of oxidase for reconstitution, the results cannot be directly compared, as the orientation of the three enzymes, and thus *pmf* and ATP generation, might vary. Nevertheless, the data show that all oxidases are capable of utilizing internal oxygen in an otherwise anoxic environment. Importantly, the various oxidases produced similar amounts ATP with catalase-produced oxygen under otherwise anaerobic conditions as under fully aerobic conditions (Figure 5A, blue bars). Normalization of the amount of ATP synthesized to the oxygen consumption rates of the reconstituted enzymes revealed that *bo*_3_ was twice as efficient as *bd* oxidases, consistent with its higher proton to electron stoichiometry (2H^+^/e^-^ *vs*. 1H^+^/e^-^; Figure S5). Although semi-quantitative, these data challenge reports of low oxygen affinity for *bo*_3_ oxidase [14,35,36]. This was supported by quinol oxidation rates of purified enzymes, which remained unaffected by the oxygen concentration (0.6 µM *vs*. 250 μM O_2_; Figure S6A). Similarly, oxygraphy , showed similar residual oxygen depletion across all oxidases (Figure S6CD). Finally, we confirmed that catalase-dependent ATP production occurs even at low (10 μM) hydrogen peroxide concentrations, typical of neutrophil defence zones [20], with all three oxidases performing similarly (Figure 5B).

**Figure 5:**
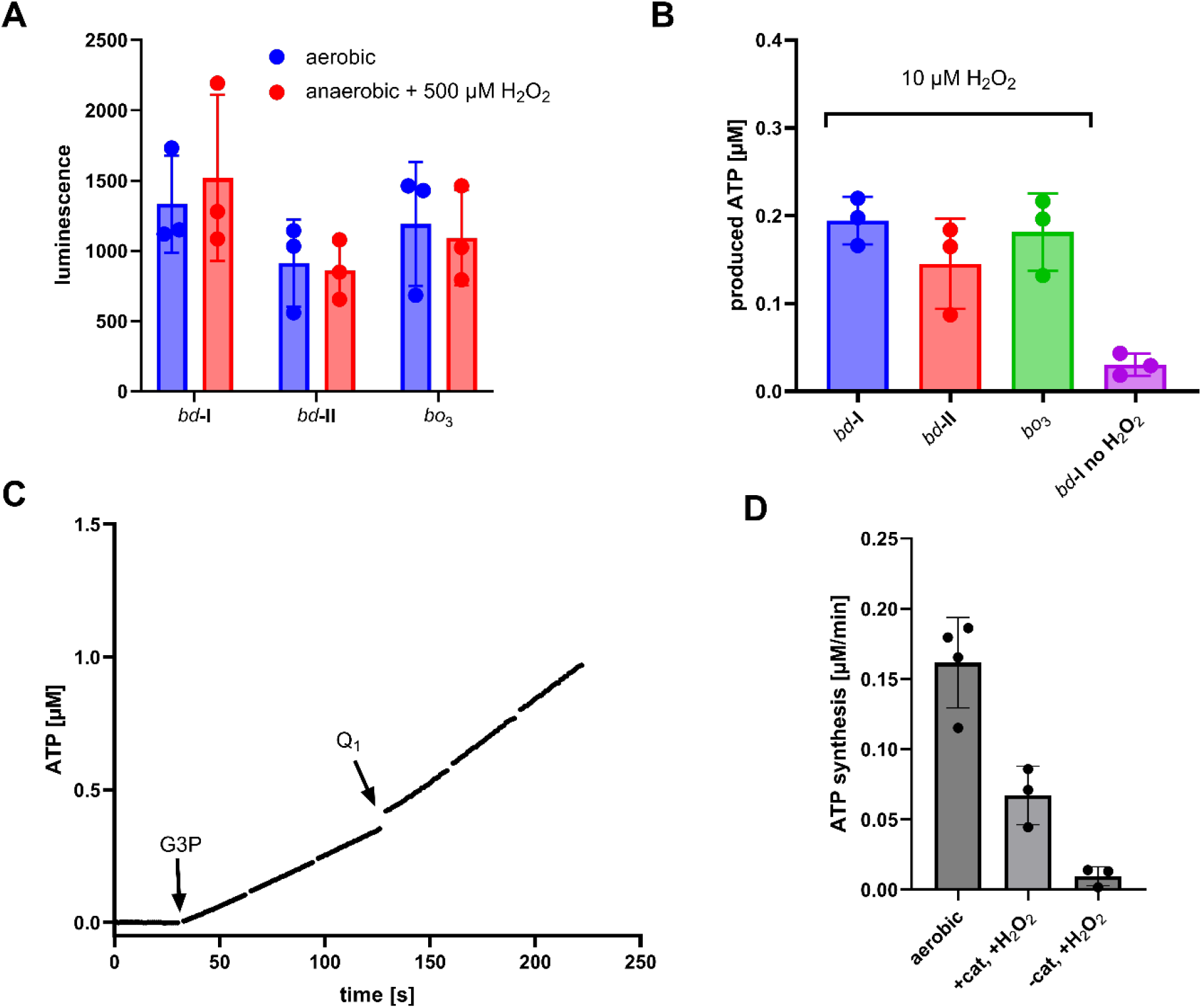
ATP synthesis of thel E¡.coli terminal oxidases and energisation by E¡.coli GlpD. A) ATP produced by *bd*-I, *bd*-II, *bo*_3_ driven ATP synthesis. Soybean lecithin proteoliposomes co-reconstituted with a terminal oxidase, ATP synthase and containing encapsulated catalase were energised by Q_1_/DTT and ATP produced after 3 minutes was measured. Aerobic measurements were started by addition of Q_1_ and anaerobic measurements by 500 µM hydrogen peroxide, respectively. B) ATP produced by lecithin proteoliposomes co-reconstituted with a terminal oxidase, ATP synthase and containing encapsulated catalase induced by an addition of 10 µM hydrogen peroxide. ATP synthesis was initiated by addition of hydrogen peroxide and run until plateau was reached. C) Aerobic ATP production of 4PC/4PE/2PG proteoliposomes containing Q_10_ co-reconstituted with *bd*-I and ATP synthase energised by GlpD. 200 nM of GlpD were added prior to the measurement and the reaction was started by addition of G3P. After 1.5 minutes short-chain Q_1_ was additionally added to the reaction. D) Rates of ATP synthesis under anaerobic conditions utilizing 4PC/4PE/2PG proteoliposomes containing Q_10_ co-reconstituted with *bd*-I, ATP synthase either with or without encapsulated catalase. Aerobic measurements were started by an addition of G3P and anaerobic measurements by hydrogen peroxide.

So far, ATP synthesis assays have relied on soluble quinones (quinone Q_1_ or Q_2_), reduced by 1,4- Dithiothreitol (DTT). However, this deviates from the natural situation where membrane- embedded, long-chain quinones are reduced by membrane-bound or membrane-associated dehydrogenases such as NADH:quinone oxidoreductase (respiratory complex I) and the alternative NADH dehydrogenase (NDH-2) or the glycerol-3-phosphate:quinone oxidoreductase GlpD. To better mimic natural conditions, we employed purified GlpD from *E. coli* to test whether anaerobic ATP synthesis is also possible with membrane-embedded long chain ubiquinone as electron mediator. GlpD is a peripheral membrane protein that associates with the membrane *via* an amphipathic alpha-helix. Rather than the natural Q_8_ found in *E. coli*, here we have used the more accessible Q_10_, which was shown to behave similarly with the *bo*_3_ oxidase [30]. We have thus co-reconstituted *bd*-I oxidase, ATP synthase into catalase-containing liposomes composed of 40% PC / 40% PE / 20% PG, a traditional mixture to mimic the *E. coli* membrane, and ubiquinone Q_10_ (4%). As depicted in Figure 5C, under aerobic conditions, glycerol 3- phosphate addition triggered immediate ATP production, whereas further addition of short- chain Q_1_ had minimal effect, suggesting efficient Q_8_ mediated electron transfer.

When the GlpD system was moved to the glove box to measure anaerobic ATP synthesis exploting catalase-produced oxygen, the system worked successfully while no ATP synthesis production was observed if catalase was lacking (Figure 5D). In comparison to the fully aerated system, ATP synthesis was less efficient under anaerobic conditions. A possible explanation might be related to the described reactive oxygen production of mitochondrial GlpD in a glycerol 3-phosphate dependent manner [37,38]. Here, GlpD may thus compete with *bd*-I oxidase for catalase-derived oxygen (producing superoxide), thereby limiting ATP synthesis.

Finally, the rate of GlpD system under fully aerobic conditions was lower compared to the DTT/Q1 system. Thermodynamically, this is to be expected as the Gibbs free energy for the GlpD reaction (ΔG°′ ≈ −44 kJ/mol) is significantly less favourable than that of the DTT/Q_1_ redox couple (ΔG°′ ≈ −71 kJ/mol).

## Conclusion

Our synthetic biology approach mimicking a minimal respiratory chain demonstrates that hydrogen peroxide can serve as a source of oxygen under otherwise anaerobic conditions and drive quinol oxidation, *pmf* generation, and ATP synthesis by employing *E. coli* terminal oxidases. Catalase efficiently decomposes hydrogen peroxide to oxygen, which is rapidly consumed by cytochrome *bd*-I, *bd*-II, or *bo*₃ oxidase, thereby detoxifying harmful reactive oxygen species into a precious substrate supporting oxidative phosphorylation under otherwise electron acceptor free conditions. Importantly, efficient utilization of the generated oxygen required catalase to be co-encapsulated within the liposomes, indicating that local production of oxygen in close proximity to the membrane-embedded oxidases is critical for effective coupling of peroxide decomposition to respiratory turnover. Oxygen availability is therefore tightly balanced between hydrogen peroxide concentration, catalase activity, and oxidase consumption by terminal oxidases.

The co-reconstitution with F_1_F_O_ ATP synthase further established that H_2_O_2_-dependent respiration can sustain proton pumping and ATP production, with yields scaling with peroxide concentration. The more physiological system using membrane-embedded ubiquinone Q_10_ and GlpD as a quinol-regenerating enzyme confirmed that endogenous substrates support this process, although at lower efficiencies. Finally, comparative analysis of oxidases showed that, in principal, all three oxidases can utilize catalase produced oxygen at hydrogen peroxide concentrations as low as 10 μM. Therefore, the observed physiological benefit restricted to *bd*-II seems to be mostly regulatory, as this pathway is governed primarily at the level of gene expression of the terminal oxidases. In accordance with their proposed mechanism, *bo*_3_ as the only proton pumping oxidase showed a two-fold higher P:O ratio.

## Supplementary information

**Supplementary Figure 1:**
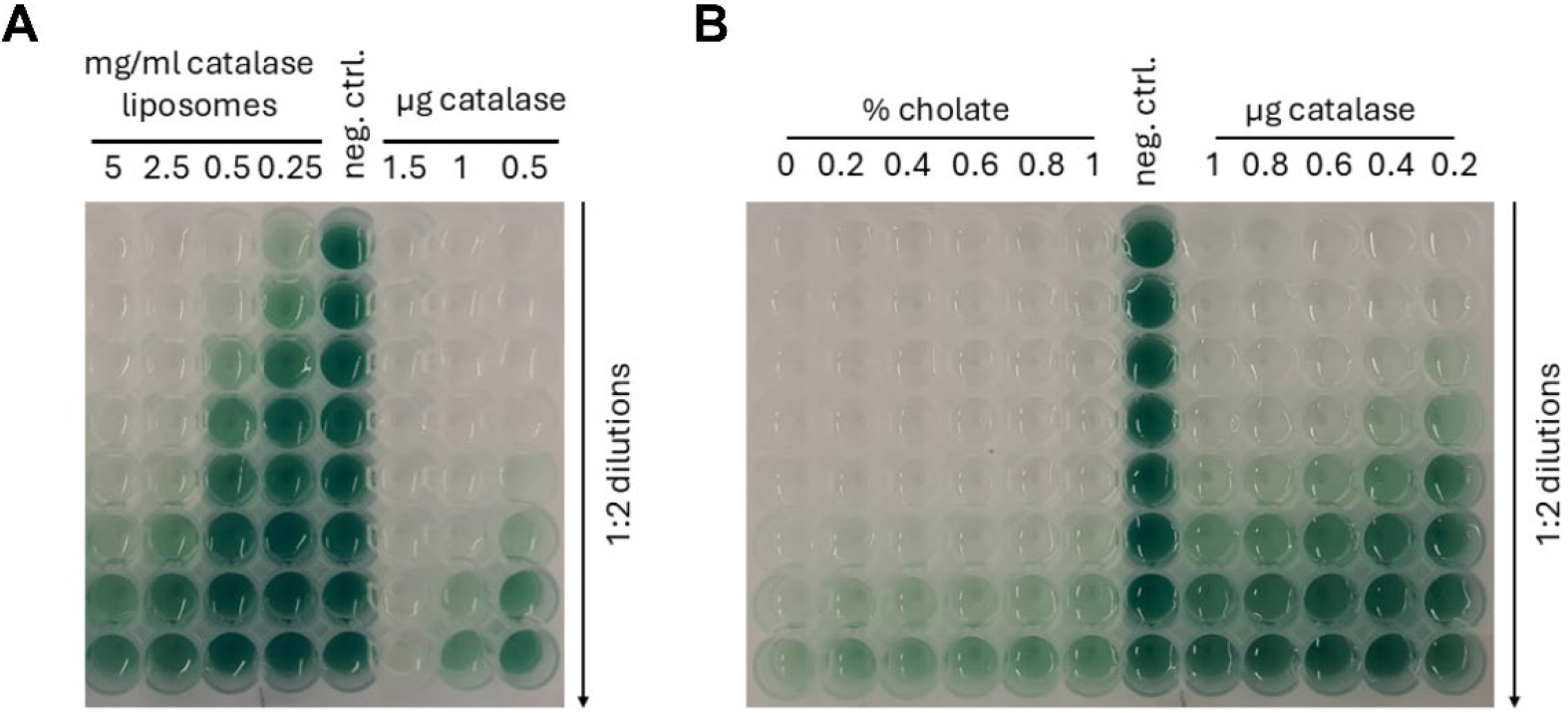
HR-POX / ABTS assays to detect catalase with sequential 1:2 dilutions. A) Catalase detection in liposomes rehydrated and freeze/thawed in presence of catalase after ultracentrifugation and standards with bovine liver catalase. B) Catalase detection in 5 mg/ml liposomes rehydrated and freeze/thawed in presence of catalase after ultracentrifugation and one hour of destabilisation with sodium cholate and detergent removal by CentriPure Z25M desalting column and standards with bovine liver catalase.

**Supplementary Figure 2:**
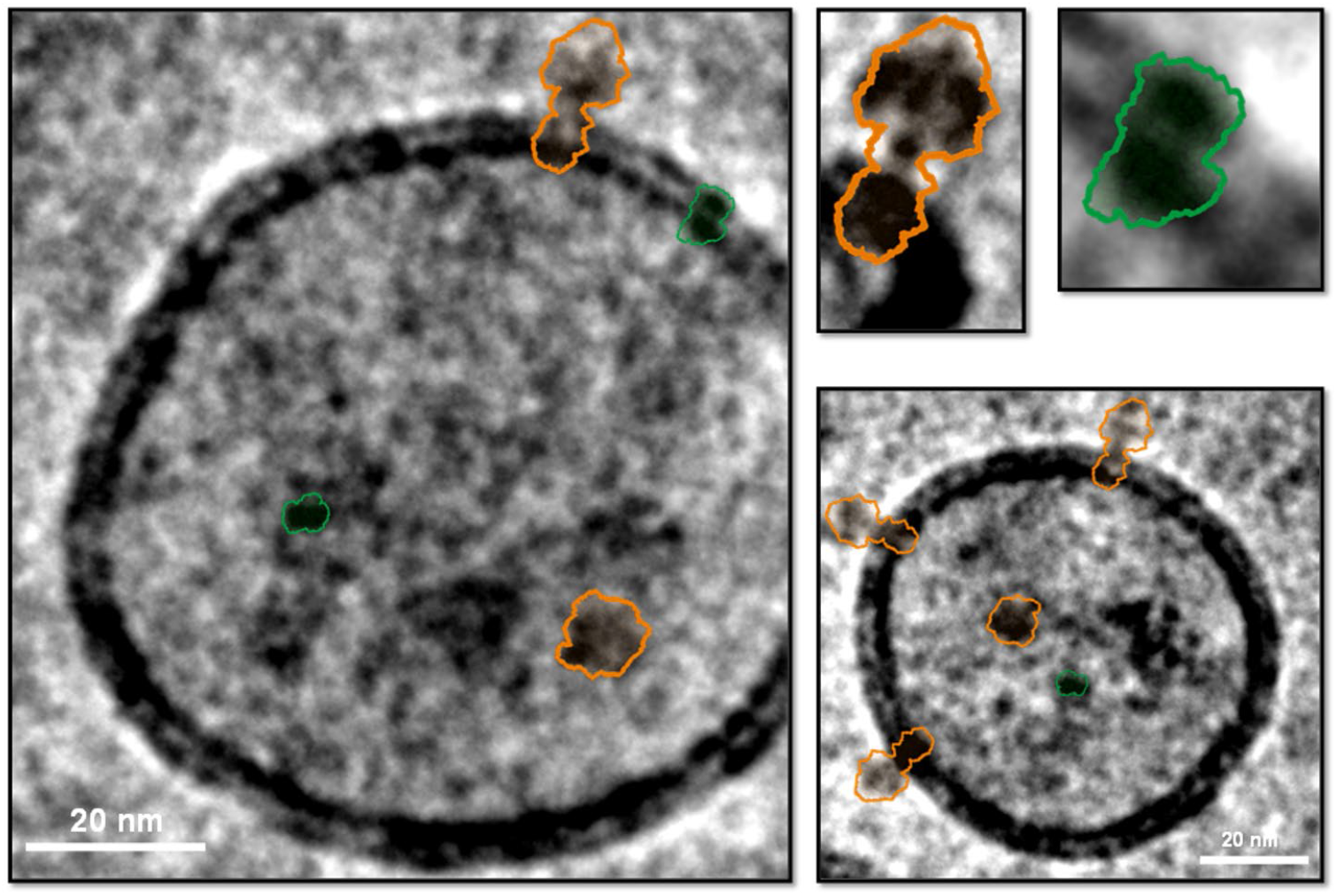
CryoEM visualization of proteoliposomes used in the study. Depicted is a selection of proteoliposomes containing reconstituted bd-II oxidase and ATP synthase from E. coli. Enzymes silhouettes to identify reconstituted proteins were derived from pdb structures of the respective enzymes. ATP synthase (orange), bd-II oxidase (green)

**Supplementary Figure 3:**
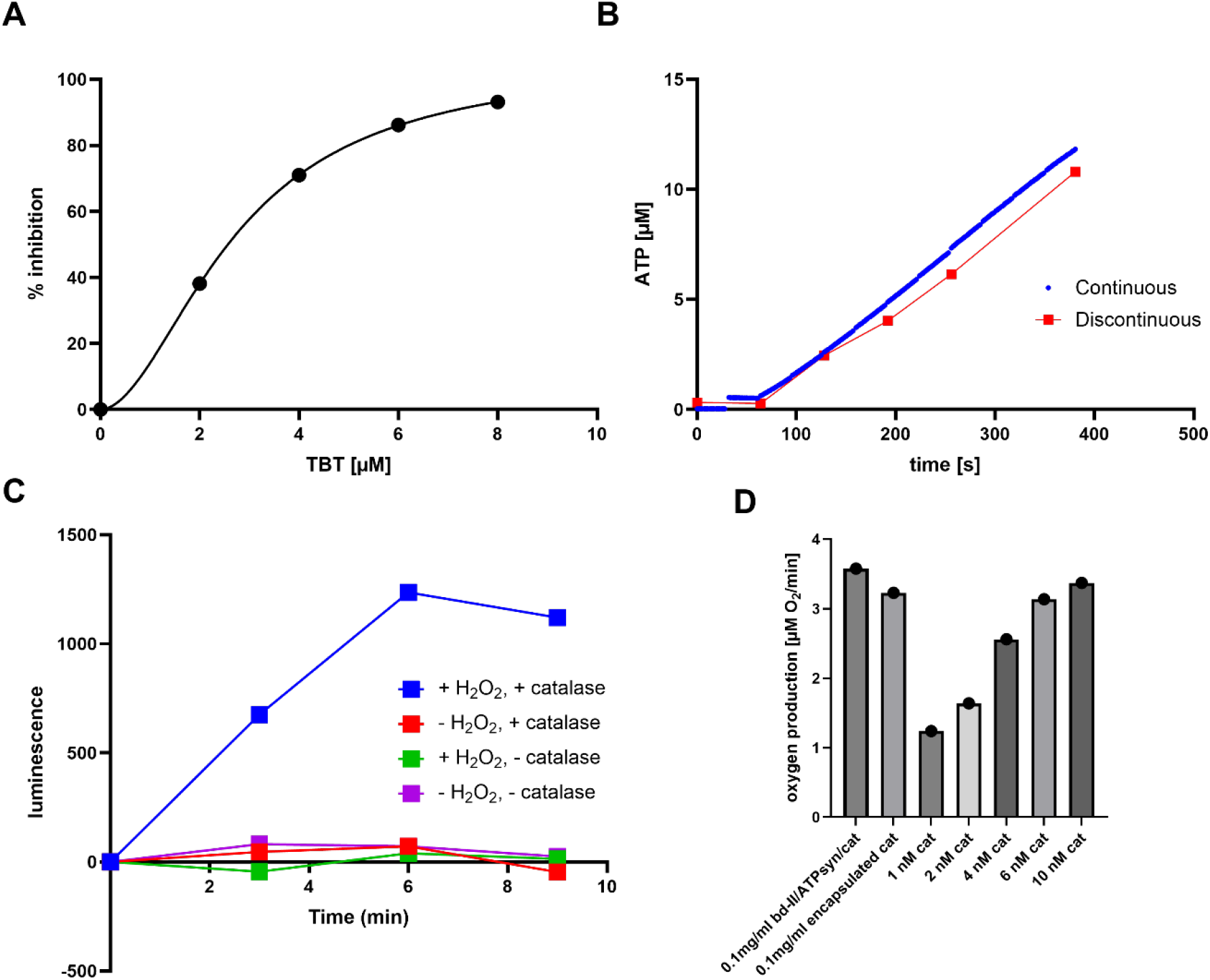
Discontinuous ATP measuring method by inhibition with TBT and controls for anaerobic hydrogen peroxide driven ATP synthesis. A) Inhibition of *bd*-I/ATP synthase ATP production by TBT. B) Comparison of continuous and discontinuous measuring methods using *bd*-I/ATP synthase proteoliposomes. C) Anaerobic ATP synthesis measured with *bd*-II and ATP synthase co-reconstituted into liposomes containing embedded catalase. D) Oxygen production rates of liposomes with encapsulated catalase and of various amounts of catalase in solution.

**Supplementary Figure 4:**
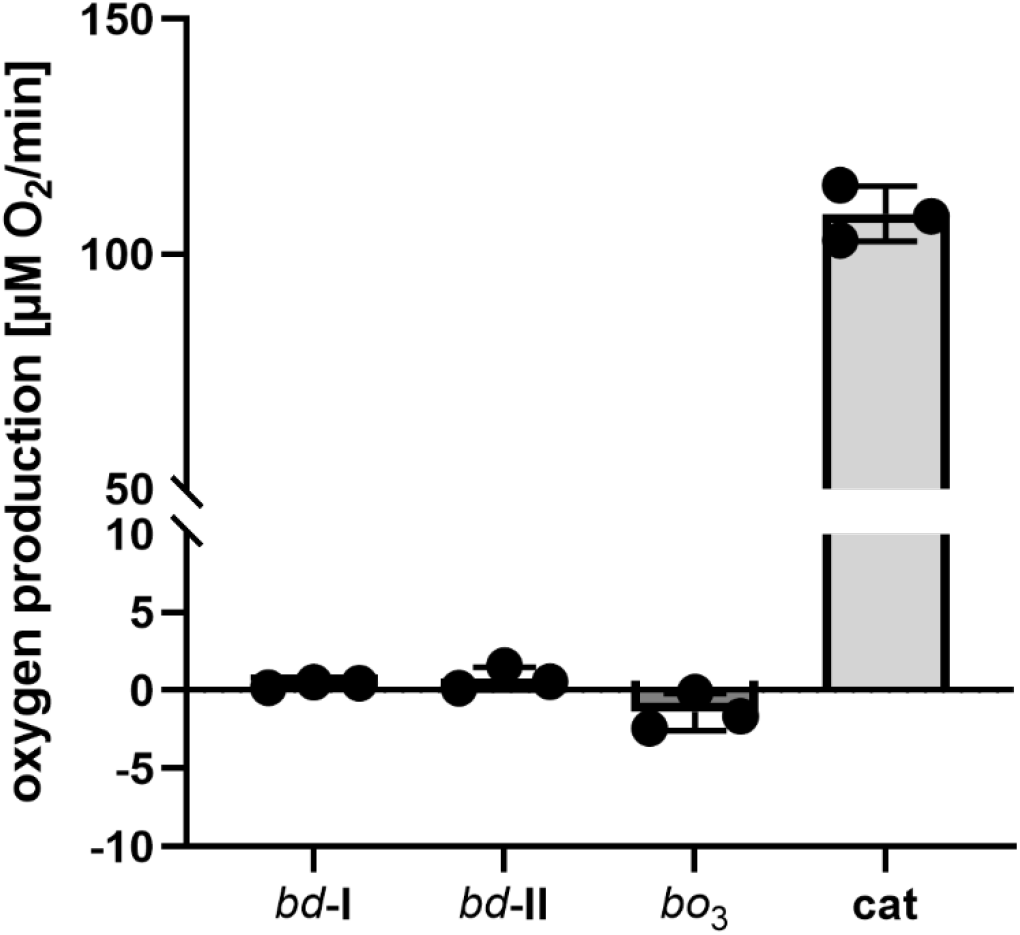
Catalase-like activity of E¡.coli terminal oxidases and bovine liver catalase. 100 nM of each enzyme was incubated with 200 µM hydrogen peroxide and the oxygen production rate was measured using a Clark-type electrode.

**Supplementary Figure 5:**
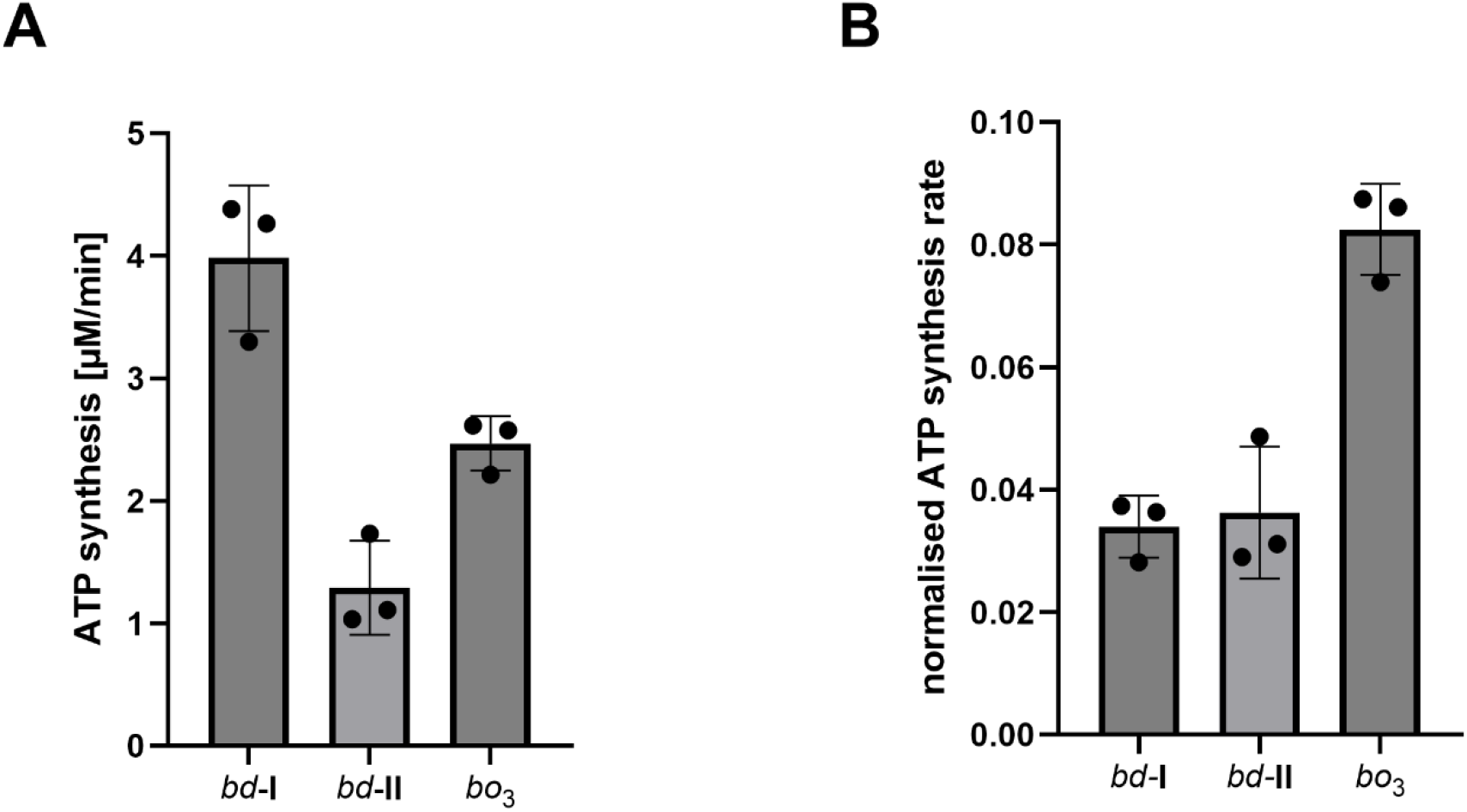
ATP synthesis of the three terminal oxidases of E¡.coli. A) ATP synthesis rate of proteoliposomes with equal amounts of *bd*-I, *bd*-II and *bo*_3_ co-reconstituted with ATP synthase. The reactions were energized energised by Q_1_/DTT and started with addition of Q_1_. B) ATP synthesis rates from A normalised to the oxygen consumption rates of liposomes energised by Q_1_/DTT.

**Supplementary Figure 6:**
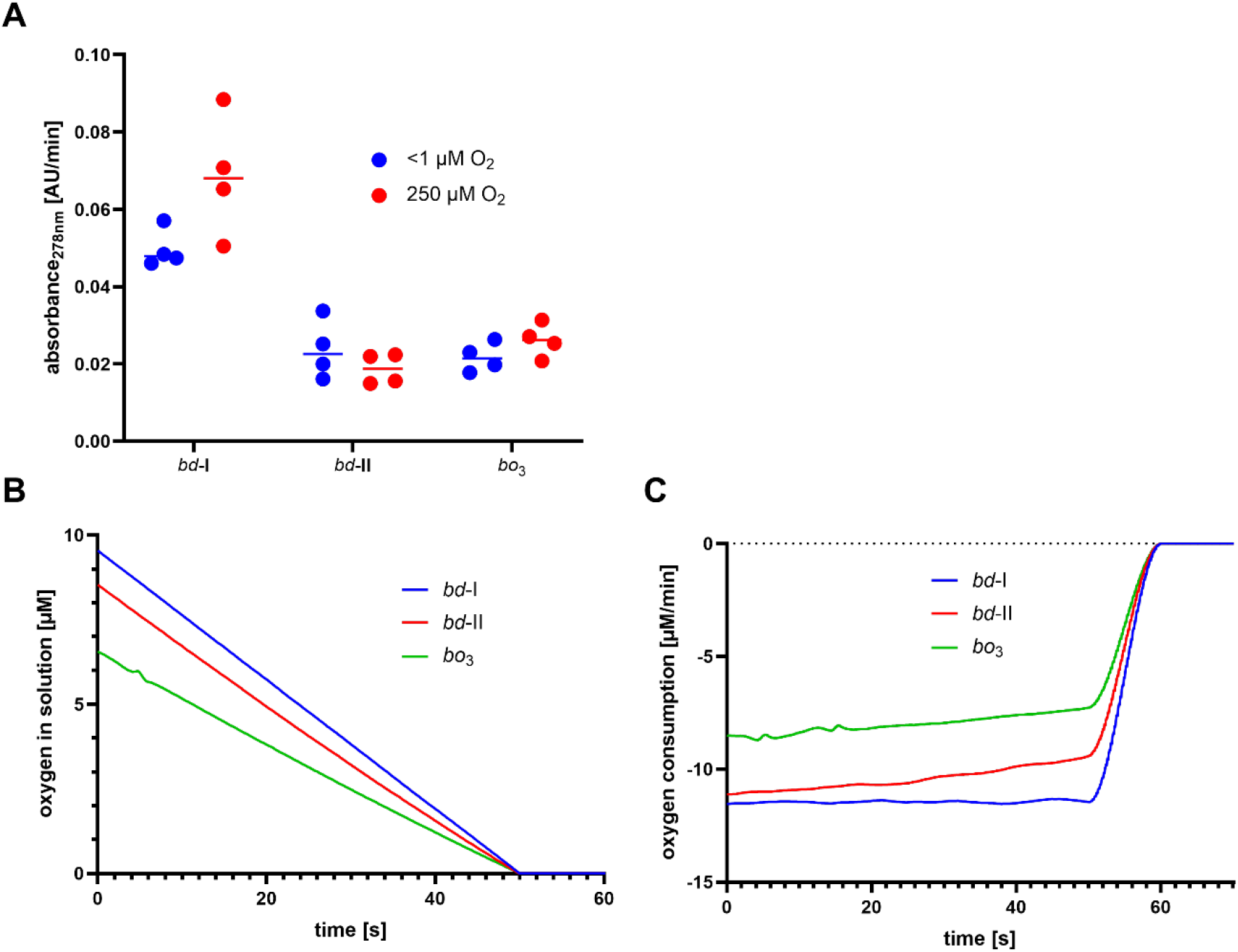
Quinol oxidation and oxygen reduction rate of E¡.coli terminal oxidases at low oxygen concentrations. A) Quinol oxidation rates of terminal oxidases at low (0.625 µM) and high (250 µM) oxygen concentrations driven by Q_1_/DTT. Low oxygen concentrations were achieved by anaerobic sample preparation with catalase present and addition of 1.25 µM of hydrogen peroxide. High oxygen concentrations were established in the same reaction by opening of the air-tight cuvette. B) Oxygen consumption of terminal oxidases at low oxygen concentrations before exhaustion of oxygen in solution driven by Q_1_/DTT. C) Oxygen consumption rates (μM/ min) extracted from the measurements shown in B.

**Supplementary Figure 7:**
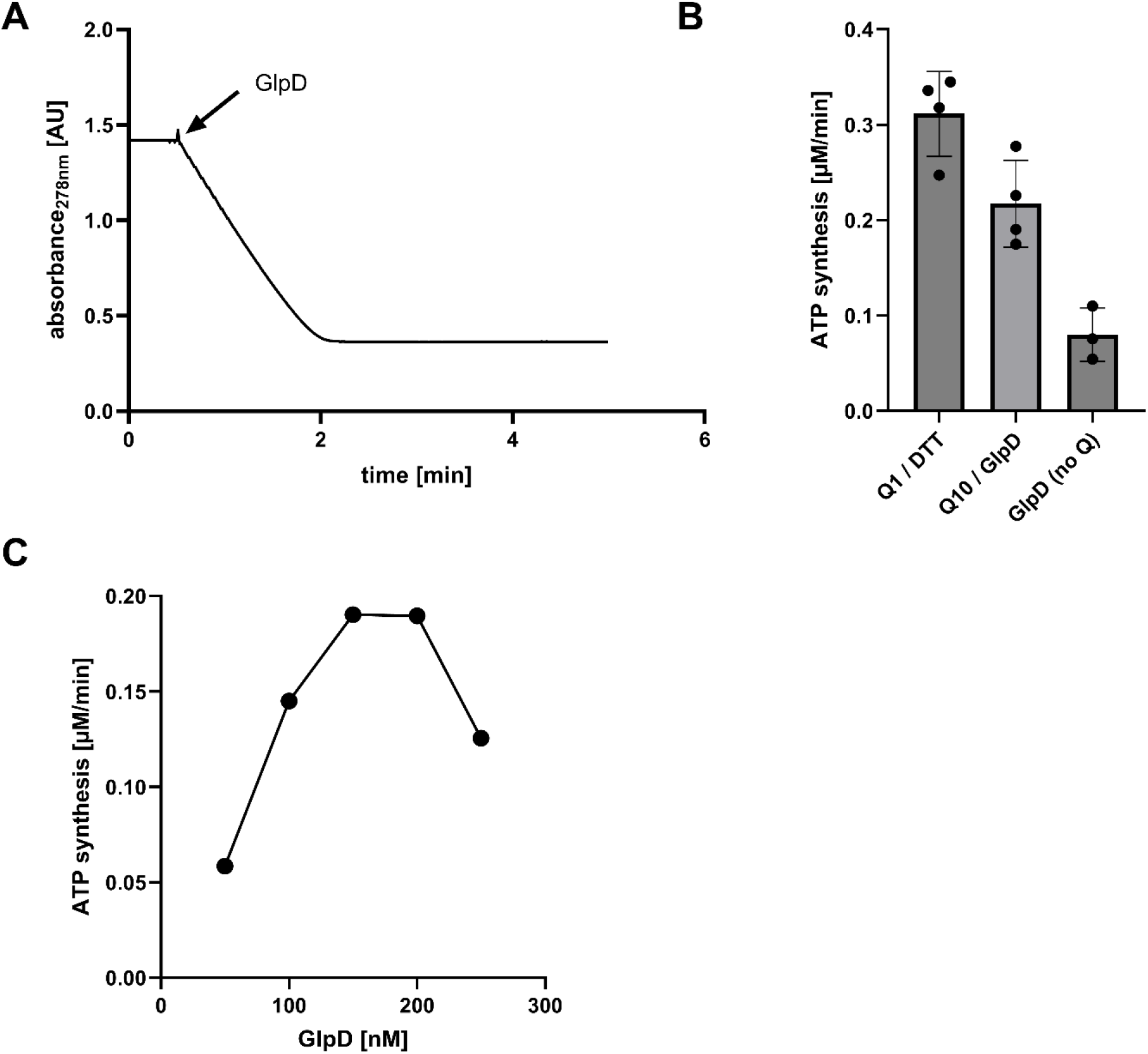
Energisation of proteoliposomes co-reconstituted with oxidase/ATP synthase by GlpD and G3P. A) Reduction of 50 µM Q_1_ by 200 nM GlpD and 6 mM G3P. B) ATP synthesis of proteoliposomes containing Q_10_ co-reconstituted with *bd*-I oxidase and ATP synthase and either energised by GlpD and G3P or with DTT and Q_1_. C) Titration of GlpD to proteoliposomes containing Q_10_ co-reconstituted with *bd*-I oxidase and ATP synthase.

## Material and Methods

### Chemicals and plasmids

If not stated otherwise chemicals were obtained from Merck / Sigma Aldrich. Ubiquinones Q_1_ and Q_2_ were synthesised as previously described [39]. For expression of GlpD the *glpD* gene from *E. coli* with a C-terminal HisTag was synthesised and cloned into a pET28a(+) expression vector by Gene Universal.

### Production and purification of proteins

Recombinant production of *E. coli bd*-I, *bd*-II, *bo*_3_, F_1_F_O_ ATP synthase, and GlpD were all done according to previous descriptions. Cytochrome *bd*-I oxidase was produced in the *E. coli* strain CBO, a C43(DE3)-based mutant lacking the cydABX and appBCX operon [40], grown in LB medium using the vector pET28b(+)cydA_his_BX. This plasmid encodes subunits A, B, and X, with a C-terminal His-tag fused to CydA to facilitate purification by immobilized metal affinity chromatography as described previously by Thessling et al [41]. Cells were broken by high-pressure homogenizer HPL6 (MAXIMATOR AG). Membranes were solubilised with 1% of lauryl maltose neopentyl glycol (LMNG, Anatrace) and applied to a Ni-NTA affinity column (Cytiva) equilibrated in buffer containing 50 mM MOPS pH 7, 500 mM NaCl, and 0.003% LMNG. The protein was subsequently eluted with an imidazole gradient ranging from 20 to 500 mM over three column volumes. Fractions corresponding to the elution peak were combined and concentrated using a 100 kDa cut-off centrifugal concentrator (Amicon Ultra, Merck). The purified enzyme was aliquoted, snap-frozen in liquid nitrogen, and stored at −80 °C until further use.

Cytochrome *bd*-II oxidase was produced in the *E. coli* strain CBO [40] using the pET28b(+) appC_his_BX plasmid and purified as previously described by Grauel et al [18]. Membrane proteins were solubilised using 1% of LMNG. The protein was purified using Ni-NTA affinity chromatography (Cytiva) equilibrated with 50 mM MOPS pH 7, 500 mM NaCl, 0.5 mM PMSF 0.003% LMNG, 50 mM imidazole. The column was washed with buffer containing 140 mM imidazole and eluted with 260 mM imidazole. Peak fractions were collected and further purified *via* size exclusion chromatography using a HiLoad 16/60 Superdex 200 pg column (GE Healthcare) equilibrated with 20 mM MOPS pH 7, 20 mM NaCl, 0.5 mM PMSF, 0.003% LMNG. Peak fractions containing *bd*-II oxidase were pooled, concentrated (50 kDa cut-off, Amicon Ultra, Merck), flash frozen in liquid nitrogen and stored at -80 °C.

Cytochrome *bo*_3_ oxidase was produced and purified as previously described by Deutschmann et al [30]. pETcyoII, encoding the *cyo* operon [42] was expressed in the *E. coli* strain C43Δcyo [43] using minimal M63 medium. Membranes were solubilised with 1% dodecyl maltoside (DDM, Glycon biochemicals GmbH) and purified *via* Ni-NTA affinity chromatography (Cytiva) equilibrated with 50 mM K2HPO4 pH 8.3, 0.05% DDM, 5 mM imidazole. The column was washed with buffer containing 35 mM imidazole and eluted with 100 mM imidazole. Peak fractions were concentrated (100 kDa cut-off, Amicon Ultra, Merck), flash frozen in liquid nitrogen and stored at -80 °C.

The *E. coli* F_1_F_O_ ATP synthase was produced from the plasmid pBWU13β-Ηis in *E. coli* DK8 cells grown in LB medium as described [44]. Membranes were solubilised with 2 % lauryl maltose neopentyl glycol (LMNG, Anatrace) and purified using prepacked Ni-NTA affinity columns (Cytiva) pre-equilibrated with 50 mM MOPS-NaOH pH 8, 100 mM NaCl, 5 mM MgCl_2_, 30 g/l sucrose, 10 % glycerol, 20 mM imidazole, 0.005 % LMNG. The column was washed with buffer containing 20 and 75 mM imidazole respectively, and the enzyme was eluted with 285 mM imidazole. The peak fractions were pooled, concentrated (100 kDa cut-off, Amicon Ultra, Merck), flash frozen in liquid nitrogen and kept at -80 °C.

GlpD from *E. coli* was produced from a pET28a(+) vector containing the *glpD* gene. Production and purification was performed according to Wright et al [45]. Briefly, the enzyme was produced in the *E. coli* RIPL expression strain grown in LB medium. Cells were broken using a high-pressure homogenizer HPL6 (MAXIMATOR AG). Isolated membranes were solubilised with 1.3 % octyl glucoside (OG, Anatrace) and the enzyme was purified using a Ni-NTA affinity column (Cytiva) equilibrated with 20 mM HEPES pH 7.4, 20 mM KCl, 200 mM NaCl and 0.05% DDM. The column was washed with buffer containing 150 mM imidazole and eluted with 400 mM. Peak fractions were concentrated (10 kDa cut-off, Amicon Ultra, Merck), frozen in liquid nitrogen and stored at -80 °C.

### Preparation of proteoliposomes

Soybean lecithin lipids (Alfa Aesar) were vigorously resuspended under nitrogen atmosphere in co-reconstitution buffer (20 mM HEPES, 50 mM KCl, 2.5 mM MgCl2, 50 g/L sucrose) for ATP synthesis or the according measuring buffer at 10 mg/mL and subjected to seven freeze/thaw cycles (LN_2_ / +29.4 °C). The resulting liposomes were made uniform in size by extrusion (21x) through a 100 nm Whatman filter (Cytiva).

Liposomes with defined headgroups were made with LIPOID E PC-S, LIPOID PG 18:1/18:1 and LIPOID E PE obtained from Lipoid. The powder was taken up in chloroform and mixed in the desired ratio (usually 4 PC / 4 PE /2 PG). A thin lipid film was obtained by rotary evaporation of chloroform and further drying overnight in a desiccator. The lipid film was taken up in co-reconstitution buffer to a concentration of 10 mg/ml and the liposomes were subjected to seven freeze/thaw cycles (LN_2_ / +29.4 °C). Before reconstitution the liposomes were extruded 21x through a 100 nm Whatman filter (Cytiva).

Encapsulation of bovine liver catalase into liposomes was performed during resuspension either of the dried lipid powder (soybean lecithin) or the lipid film (lipoid) by addition of 50 µM of catalase to the buffer. During the freeze-thaw cycles the lipid concentration was kept at 50 mg/ml and only before extrusion the liposomes were diluted to 10 mg/ml.

Detergent mediated (co-)reconstitution of terminal oxidases and F_1_F_O_ ATP synthase was performed with 0.4% of sodium cholate to destabilise the preformed liposomes. The desired enzymes were added, usually 10-20 oxidases and 5-10 ATP synthases per vesicle, and the mixture was incubated for an hour on ice with occasional gentle flicking. The sodium cholate was removed using small desalting columns (CentriPure Z25M, emp Biotech). For experiments with catalase the liposomes were collected (200’000 xg, 1 h, 4 °C) to remove external catalase.

4% (w/w) of coenzyme Q_10_ were incorporated into lipoid liposomes during mixing of the chloroform stocks. In order to efficiently incorporate the very hydrophobic electron carrier the buffer for lipid film rehydration was preheated to 50 °C. Thawing during the freeze/thaw cycles was performed done at 50 instead of 29.4 °C and also extrusion was done at 50 °C using a heating block (Avanti Polar Lipids).

### Detection of catalase

The presence of active catalase was assessed using a horseradish peroxidase / 2,2’-azino-bis(3- ethylbenzothiazoline-6-sulfonic acid (HR-POX / ABTS) assay. The wells were coated with 3% milk powder solution for 5 minutes. Samples were diluted in 20 mM tris-HCl pH 7.75 and serially diluted in 8 steps (1:2 dilutions). After 15 minutes incubation with 0.0005% H_2_O_2_ solution, 0.016 mg/ml HR-POX and 0.3 mg/ml ABTS were added, leading to green colour when peroxide is still present. For detection of catalase in liposomes the samples were first diluted to 8 ml (usually from 100 – 200 µl) and collected *via* ultracentrifugation (200’000 xg, 1h, 4°C).

### Determination of oxygen concentration in solution

The titration of hydrogen peroxide dependent oxygen production of catalase was measured using a Clark type electrode (Oxygraph+, Hansatech Instruments). Oxygen production of 200 nM of bovine liver catalase was measured in 20 mM MOPS-BTP pH 7, 50 mM KCl, 2.5 mM MgCl_2_ and was repeated in presence of 15 µM Q_1_ and 4 mM DTT. The reaction was started with the addition of catalase and run until a plateau was reached. Catalase activity in liposomes was measured in the same buffer with 0.1 mg/ml final lipid concentration (4 PC / 4 PE /2 PG). The reaction was started by addition of 1 mM hydrogen peroxide.

Diffusion of catalase derived oxygen into the bulk solution was measured with proteoliposomes with bovine liver catalase encapsulated and *bd*-II oxidase reconstituted into soybean lecithin liposomes. 20 mM MOPS-BTP pH 7, 50 mM KCl, 2.5 mM MgCl_2_ was bubbled for 30 min with argon (Carbagas) and transferred into the Oxygraph preflushed with argon. Proteoliposomes were added (0.1 mg/ml final) and for measurements with oxidase 15 µM Q_1_ and 4 mM DTT were also added. Upon establishment of a stable baseline H_2_O_2_ was added and oxygen concentration in solution was monitored either until a plateau was reached or all produced oxygen was used up by *bd*-II oxidase.

Oxygen consumption of terminal oxidases was measured in 20 mM MOPS pH 7, 20 mM NaCl, 0.05% DDM, 4 mM DTT, 15 µM Q_1_ using the Oxygraph+ (Hansatech Instruments). For normalisation of ATP synthesis rates oxygen consumption was measured with proteoliposomes also used for the corresponding ATP synthesis measurements (2 mg/ml) and without detergent. For measurement of oxygen consumption rates at low oxygen levels the amount of oxidases was adjusted to achieve similar consumption rates.

### Quinol oxidation by terminal oxidases

Q_2_ was pre-reduced as previously described [24] in argon saturated EtOH by stepwise addition of sodium borohydride. After incubation on ice (15 min) excess sodium borohydride was removed by addition of HCl and white precipitation was cleared by centrifugation (10’000 xg, 10 min, 4 °C).

For anaerobic measurements the reaction mixture containing 20 mM MOPS pH 7, 20 mM NaCl, 0.003% LMNG, 45 nM bovine liver catalase and 133 µM pre-reduced Q_2_ was bubbled for 30 minutes with argon (Carbagas) in an air-tight cuvette (Hellma Analytics) or stirred in an anaerobic glovebox for at least an hour. Quinol oxidation was followed at 275 nm using a Cary 60 UV-vis spectrometer (Agilent Technologies). At the indicated marks, 25 nM of *bd*-II oxidase and hydrogen peroxide were added and the reaction was run until a plateau was reached. Controls were done without argon bubbling or no addition of *bd*-II oxidase. Measurements with proteoliposomes were done with catalase encapsulated and *bd*-II reconstituted into soybean lecithin liposomes. Prior to measurement the proteoliposomes and buffer were stirred in an anaerobic hood for one hour. Reaction was initiated by an addition of 0.1 mg/ml proteoliposomes and 200 µM H_2_O_2_ at the indicated timepoints.

Quinol oxidation at low oxygen concentrations was measured in 20 mM NaCl, 0.003% LMNG with 30 µM pre-reduced Q_1_ and 50 nM bovine liver catalase. All components were incubated and mixed in an anaerobic glovebox and measured in an airtight cuvette (Hellma Analytics). The reactions were started by addition of 1.25 µM of H_2_O_2_, resulting in in a maximal oxygen concentration of 0.625 µM oxygen.

### Quinone reduction by GlpD

Enzymatic activity of *E. coli* GlpD was assessed by spectroscopic measurement of quinone reduction at 278 nm using a Cary 60 UV-vis spectrometer (Agilent Technologies). 200 nM GlpD were mixed with 20 mM HEPES pH 7.4, 20 mM NaCl, 20 mM KCl, 50 µM Q_1_ and the reaction was started by addition of 6 mM of glycerol-3-phosphate (G3P).

### Determination of ATP synthesis rates

#### Aerobic continuous ATP synthesis

ATP production was always performed with F_1_F_O_ ATP synthase co-reconstituted with a terminal oxidase (*bd*-I, *bd*-II or *bo*_3_) at a proteoliposome concentration of 0.1 mg/ml (if not indicated otherwise) in 20 mM tris-PO_4_ pH 7.4, 50 mM KCl, 2.5 mM MgCl_2_, 200 mM ADP, 0.4 mg/ml luciferin/luciferase mix. ATP concentration was monitored using an ATP Bioluminescence Assay Kit CLS II (Roche) and luminescence was measured with a GloMax 20/20 Luminometer (Promega). ATP concentrations were quantified by addition of a known amount of ATP (usually 0.1 µM) prior to the measurement.

#### Anaerobic discontinuous ATP synthesis

For discontinuous ATP measurements the reaction and luminescent measurements were performed in two different vessels. Reactions were run in 20 mM tris-PO_4_, 50 mM KCl, 2.5 mM MgCl_2_, 200 mM ADP with 0.1 mg/ml proteoliposomes containing F_1_F_O_ ATP synthase and an oxidase. During the reaction aliquots were withdrawn and diluted in 20 mM MOPS-BTP pH 7, 50 mM KCl, 10 µM TBT, 0.4 mg/ml luciferin/luciferase mix (ATP Bioluminescence Assay Kit CLS II, Roche), stopping the reaction and measured with a GloMax 20/20 Luminometer (Promega). Luminescence was calibrated by addition of 0.1 µM ATP.

For anaerobic measurements proteoliposomes with encapsulated bovine liver catalase were used. All buffer and reaction components were first incubated at least for two hours and then mixed in an anaerobic glovebox, whereafter the mixture was stirred for another 20 minutes in absence of oxygen. ATP synthesis was started by hydrogen peroxide addition and run in airtight cuvettes (Hellma Analytics), samples were withdrawn with gastight syringes (Hamilton) and immediately mixed with 20 mM MOPS-BTP pH 7, 50 mM KCl, 10 µM TBT, 0.4 mg/ml luciferin/luciferase mix (ATP Bioluminescence Assay Kit CLS II, Roche) for measurement with a GloMax®20/20 Luminometer (Promega).

#### Energisation of pmf dependent ATP synthesis

If not otherwise indicated, ATP synthesis energised by Q_1_/DTT was measured in soybean lecithin liposomes. The buffer was supplemented with 4 mM DTT and after recording of the baseline the reaction was started by addition of 15 µM Q_1_. Reactions energised by GlpD/glycerol-3-phosphate (G3P) were performed in lipoid proteoliposomes (4 PC / 4 PE /2 PG) with incorporated Q_10_. Prior to the measurement 200 nM of GlpD was added and the reaction was started with 5 mM of G3P.

